# Bovine Lactoferrin Activity Against Chikungunya and Zika Viruses

**DOI:** 10.1101/071571

**Authors:** Carlos A. M. Carvalho, Samir M. M. Casseb, Rafael B. Gonçalves, Eliana V. P. Silva, Andre M. O. Gomes, Pedro F. C. Vasconcelos

## Abstract

Chikungunya (CHIKV) and Zika (ZIKV) viruses are two arboviruses which have recently broken their sylvatic isolation and gone into a rampant spreading among humans in some urban areas of the world, specially in Latin America. Given the huge burden that Chikungunya and Zika fevers impose to public health in the affected countries and the lack of effective interventions against them, the aim of this work was to evaluate the antiviral potential of bovine lactoferrin (bLf) – an iron-binding glycoprotein with broad-spectrum antimicrobial properties – in both CHIKV and ZIKV infections. The general antiviral activity of bLf was assessed by plaque assays, and the inhibitory effects of the protein on specific stages of virus infecion was evaluated by immunofluorescence and nucleic acid quantification assays. Our data show that bLf exerts a dose-dependent strong inhibitory effect on the infection of Vero cells by the aforementioned arboviruses, reducing their infection efficiency in up to nearly 80%, with no significant cytotoxicity, and such antiviral activity occurs at the levels of binding and replication of the virus particles. Taken together, these findings reveal that bLf antimicrobial properties are extendable to CHIKV and ZIKV, underlining a generic inhibition mechanism that can be explored to develop a potential strategy against their infections.

## Introduction

Over the past few years, the world has witnessed epidemics of human infections caused by two old acquaintance, yet still obscure, arboviruses: Chikungunya (CHIKV) and Zika (ZIKV) viruses. While CHIKV is a member of the *Alphavirus* genus in the *Togaviridae* family, first isolated in Tanzania, in 1952 [1, 2], ZIKV is a member of the *Flavivirus* genus in the *Flaviviridae* family, first isolated in Uganda, in 1947 [3, 4]. These viruses are mainly transmitted by mosquitoes belonging to the *Aedes* genus, and are the etiological agent of dengue-like febrile illnesses that show a range of superimposing non-specific signals and symptoms, which may constitute a syndromic framework [5]. Chikungunya fever is frequently associated with a high prevalence of chronic arthralgia and ZIKV may be associated with congenital microcephaly [6]. As for other arbovirus diseases, no effective antiviral intervention is hitherto available for cases of Chikungunya or Zika fevers [7].

In the urge for a means to halt the infection by multiple viruses, broad-spectrum drugs from nature may provide valuable hints, since the life cycle of different virus species share common cellular factors and pathways [8]. Among these drugs, lactoferrin (Lf) – an iron-binding globular glycoprotein of about 700 amino acid residues belonging to the transferrin family [9] – is noteworthy. First isolated from bovine (bLf) and human (hLf) milk in 1960 [10, 11], Lf is also found in various mucosal secretions, such as tears, saliva and seminal/vaginal fluids, and in the secondary granules of mature neutrophils [12, 13], playing an important role in the primary defense against a broad spectrum of pathogenic microorganisms, including bacteria, protozoa, fungi and many naked and enveloped viruses [14].

The aim of this study was to evaluate the antiviral potential of bLf in CHIKV and ZIKV infections as a way to gather clues for the development of efficient therapeutic interventions. Using plaque, immunofluorescence, and nucleic acid quantification assays, we tested the ability of bLf to inhibit the infection of Vero cells by these viruses and attempted to determine the stages of the infection cycle at which the protein imposes its antiviral effects. Our results demonstrate that bLf exerts a non-cytotoxic strong inhibitory effect on both CHIKV and ZIKV infections at the levels of virus binding and replication, further extending its antimicrobial spectrum to such emerging arboviruses and identifying common events in their life cycles that are liable to inhibition.

## Materials and Methods

### Cell Culture

African green monkey kidney (Vero) cells (American Type Culture Collection, Manassas, USA) were cultured as monolayers in 25-cm^2^ ventilated flasks (Nest, Wuxi, China) at 37 °C in a humidified atmosphere with 5% CO2 in 199 medium (Cultilab, Campinas, Brazil) supplemented with 5% fetal bovine serum (LGC Biotecnologia, Cotia, Brazil) and 1% antibiotic antimycotic solution consisting of 10,000 U penicillin, 10 mg streptomycin, and 25 µg amphotericin B per mL (Sigma-Aldrich, St. Louis, USA).

### Virus Propagation, Clarification, and Titration

Vero cells were grown to quasi-confluence in 75-cm^2^ ventilated flasks (Nest) and then infected with Brazilian strains of CHIKV (BeH807658) or ZIKV (BeH815744) under a multiplicity of infection (MOI) of 0.1 plaque-forming unit (PFU)/cell for 48 or 96 h at 37 °C, respectively. After virus propagation, the culture medium was collected and cleared of cell debris by centrifuging at 8,872 x g for 20 min at 4 °C. The supernatant was collected, aliquoted and stored as clarified virions at -70 °C until further use. Infectious titers of virus samples were determined by plaque assay in Vero cells as described elsewhere [15].

### BLf Preparation

Encapsulated apolactoferrin from bovine whey (Life Extension, Fort Lauderdale, USA) was prepared as previously described [16]. Briefly, the protein contained in the capsules was dissolved to a concentration of 100 mg/mL in phosphate-buffered saline (PBS) and centrifuged at 4,991 x g for 5 min at 4 °C to remove the cellulose excipient. The supernatant was passed through a 0.22-μm syringe-driven filter unit (Jet Biofil, Guangzhou, China), aliquoted and stored as a stock solution at -20 °C until further use. This procedure eliminated all excipients described in the formulation provided by the manufacturer. Protein purity was 95%, as stated by the manufacturer and confirmed by sodium dodecyl sulfate-polyacrylamide gel electrophoresis (SDS-PAGE).

### Cytotoxicity Assay

Vero cell monolayers seeded in 96-well plates (Nest) were incubated with different concentrations of bLf at 37 °C for 48 h and then assayed for the cleavage of the fluorogenic, cell-permeant, peptide substrate glycylphenylalanyl-aminofluorocoumarin (GF-AFC), provided in the CellTiter-Fluor Cell Viability Assay (Promega, Fitchburg, USA), by a conserved and constitutive protease within live cells, on the GloMax-Multi+ Microplate Multimode Reader (Promega).

### Antiviral Assays

All general assays for assessing the activity of bLf against CHIKV or ZIKV infection were conducted in Vero cell monolayers seeded in 12-well plates (Nest). The dose-response activity of bLf was evaluated by incubating cells with the indicated concentrations of the protein at 37 °C throughout the course of infection, including immediately before (for 1 h), during (for 1 h), and immediately after (for 48 or 96 h, respectively) virus addition, at a density of 100 PFU/well. For the time-of-addition assays, the experimental procedure was the same, except bLf was present at a single concentration (1.0 mg/mL) separately for each stage of infection. At 48 h post-infection for CHIKV or 96 h post-infection for ZIKV, cells were stained with 0.1% crystal violet, and the virus plaques were counted to determine the efficiency of infection. Alternatively, virus samples were pretreated with 1.0 mg/mL bLf for 1 h at 37 °C, diluted to reduce the concentration of bLf far below the minimum inhibitory concentration, and titrated by plaque assay in Vero cells.

### Indirect Immunofluorescence Assay (iIFA)

Vero cells monolayers seeded in 4-well Lab-Tek II Chamber Slide Systems (Nunc, Roskilde, Denmark) were incubated with 1.0 mg/mL bLf (∼10^10^ protein molecules/cell) for 15 min at 4 °C, washed to remove unbound protein molecules, and then incubated with CHIKV or ZIKV under an MOI of 1 PFU/cell for another 15 min at 4 °C, being afterwards washed again to remove unbound virus particles and incubated at 37 °C to allow for infection progress. At 24 h post-infection for CHIKV or 48 h post-infection for ZIKV, cells were fixed with 3.7% formaldehyde for 15 min, permeabilized with 0.25% Triton X-100 for another 15 min, and blocked with 3% bovine serum albumin (BSA) for 1 h. Homemade anti-CHIKV or anti-ZIKV primary mouse polyclonal antibodies, obtained from the ascitic fluid of Swiss mice after intraperitoneal inoculations of live viruses as described elsewhere [17], were incubated with the cells at a dilution of 1:20 for 1 h, followed by incubation with FITC-conjugated anti-mouse IgG secondary goat polyclonal antibodies (Sigma-Aldrich) at a dilution of 1:500 for another 1 h. Cellular nuclei were stained by incubation with 1 µM Hoechst 33342 (Molecular Probes, Eugene, USA) for 10 min. All steps were carried out at room temperature and cells were washed with PBS after every incubation. Images were acquired on the BX51 System Microscope (Olympus, Tokyo, Japan) coupled to an X-Cite 120Q excitation light source (EXFO, Quebec, Canada) and processed using ImageJ 1.48 software (National Institutes of Health, Bethesda, USA).

### Quantitative Reverse Transcripton-Polymerase Chain Reaction (qRT-PCR)

Vero cell monolayers seeded in 6-well plates (Nest) were incubated with CHIKV or ZIKV under an MOI of 0.1 PFU/cell for 1 h at 37 °C, washed to remove unbound virus particles, and then incubated with 1.0 mg/mL bLf (∼10^10^ protein molecules/cell) for another 1 h at 37 °C, being afterwards washed again to remove unbound protein molecules and incubated at 37 °C to allow for infection progress. At 24 h post-infection for CHIKV or 48 h post-infection for ZIKV, culture media were harvested, clarified as described above, and subjected to RNA isolation using the Maxwell 16 LEV simplyRNA Cells Kit (Promega) on the Maxwell 16 Instrument (Promega). Reverse transcription and DNA amplification were performed on the 7500 Real-Time PCR System (Applied Biosystems, Foster City, USA) using the SuperScript III Platinum One-Step qRT-PCR Kit with ROX (Invitrogen, Carlsbad, USA) in addition to previously described primer/probe sets (Integrated DNA Technologies, Coralville, USA) against defined sequences in CHIKV (forward primer = 5’ – A A A G G G C A A A C T C A G C T T C A C – 3’; reverse primer = 5’ – G C C T G G G C T C A T C G T T A T T C – 3’; FAM-labeled primer = 5’ – C G C T G T G A T A C A G T G G T T T C G T G T G – 3’) or ZIKV (forward primer=5’ – C C G C T G C C C A A C A C A A G – 3’; reverse primer=5’ – C C A C T A A C G T T C T T T T G C A G A C A T – 3’; FAM-labeled primer=5’ – A G C C T A C C T T G A C A A G C A G T C A G A C A C T C A A – 3’) genomes [18, 19].

### Statistical Analyses

Statistical analyses were performed using one/two-way ANOVA with Tukey’s post-test for multiple comparisons on Prism 6 software (GraphPad Software, San Diego, USA). Data were expressed as mean ± standard deviation (SD), and P-values less than 0.05 were considered statistically significant.

## Results

### Effect of BLf on Cell Viability

In order to assess whether bLf treatment would lead to toxic effects in Vero cells, a viability assay was carried out after incubating the cells with a range of bLf concentrations for 48 h at 37 °C. All concentrations tested showed no significant cytotoxicity – even at the highest bLf concentration tested (1.0 mg/mL), Vero cell viability was still retained (Fig 1).

**Fig 1.**
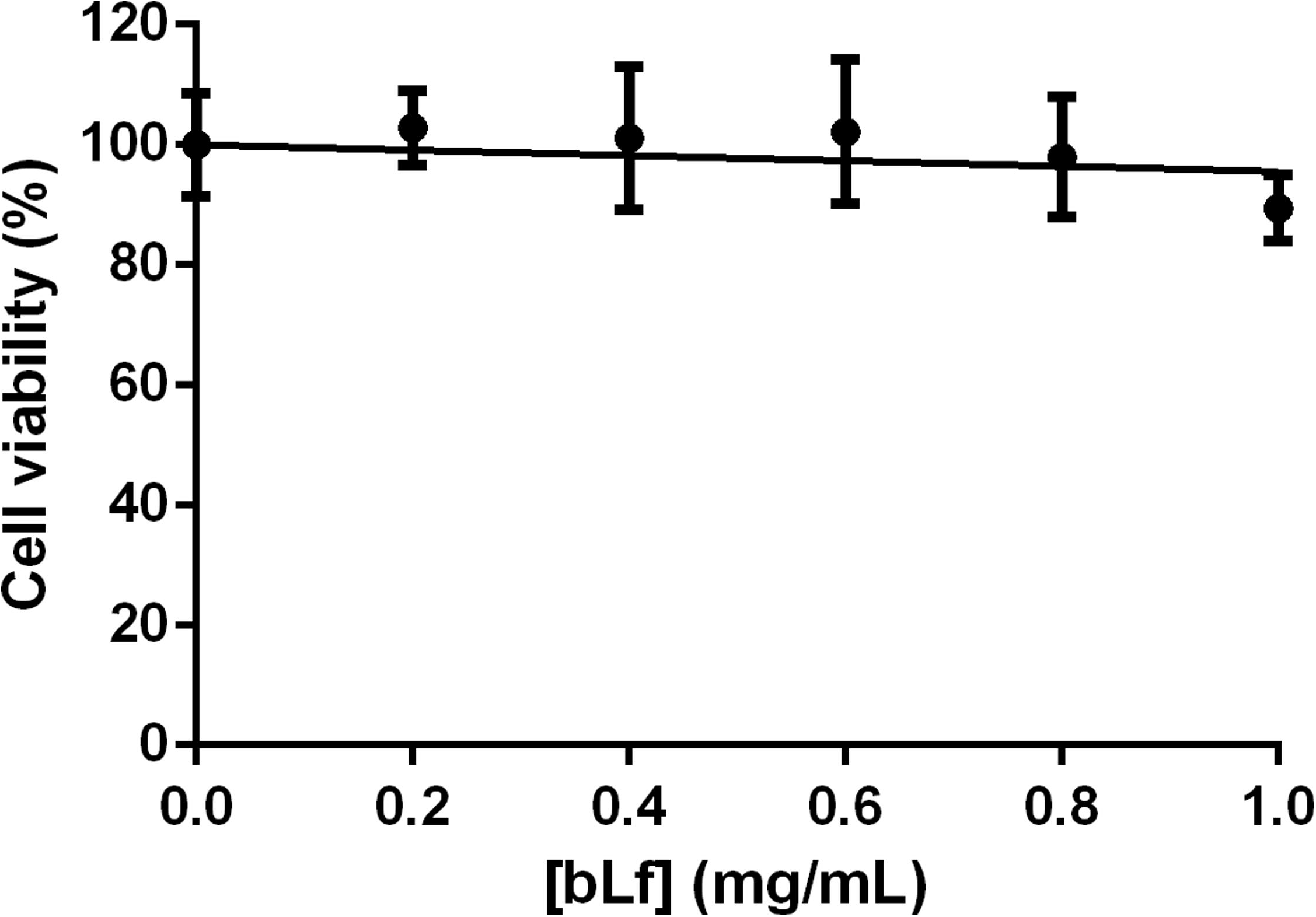
Effect of bLf on cell viability. Monolayers of Vero cells were treated with the indicated concentrations of bLf for 48 h at 37 °C and then subjected to a GF-AFC cleavage assay to determine cell viability. Data were obtained from 6 experiments and plotted as mean ± SD along with their linear regression. All differences compared to the control were not significant (P≥0.05).

### Dose-Dependent Inhibitory Effect of BLf on CHIKV or ZIKV Infection

Given the lack of cytotoxicity in the range of 0.2 to 1.0 mg/mL, bLf was assayed for its antiviral potential in CHIKV or ZIKV infection in Vero cells under these concentrations. In such assay, bLf was incubated along the whole infection procedure, including a pretreatment step for 1 h at 37 °C, and its ability to promote plaque number reduction was then tested. BLf showed a remarkable dosedependent antiviral activity, similarly preventing CHIKV or ZIKV infection by nearly 80% at a concentration of 1.0 mg/mL (Fig 2). However, the half maximal inhibitory concentration (IC_50_) of bLf was 0.2 ± 0.005 mg/mL for CHIKV and 0.4 ± 0.006 mg/mL for ZIKV.

**Fig 2.**
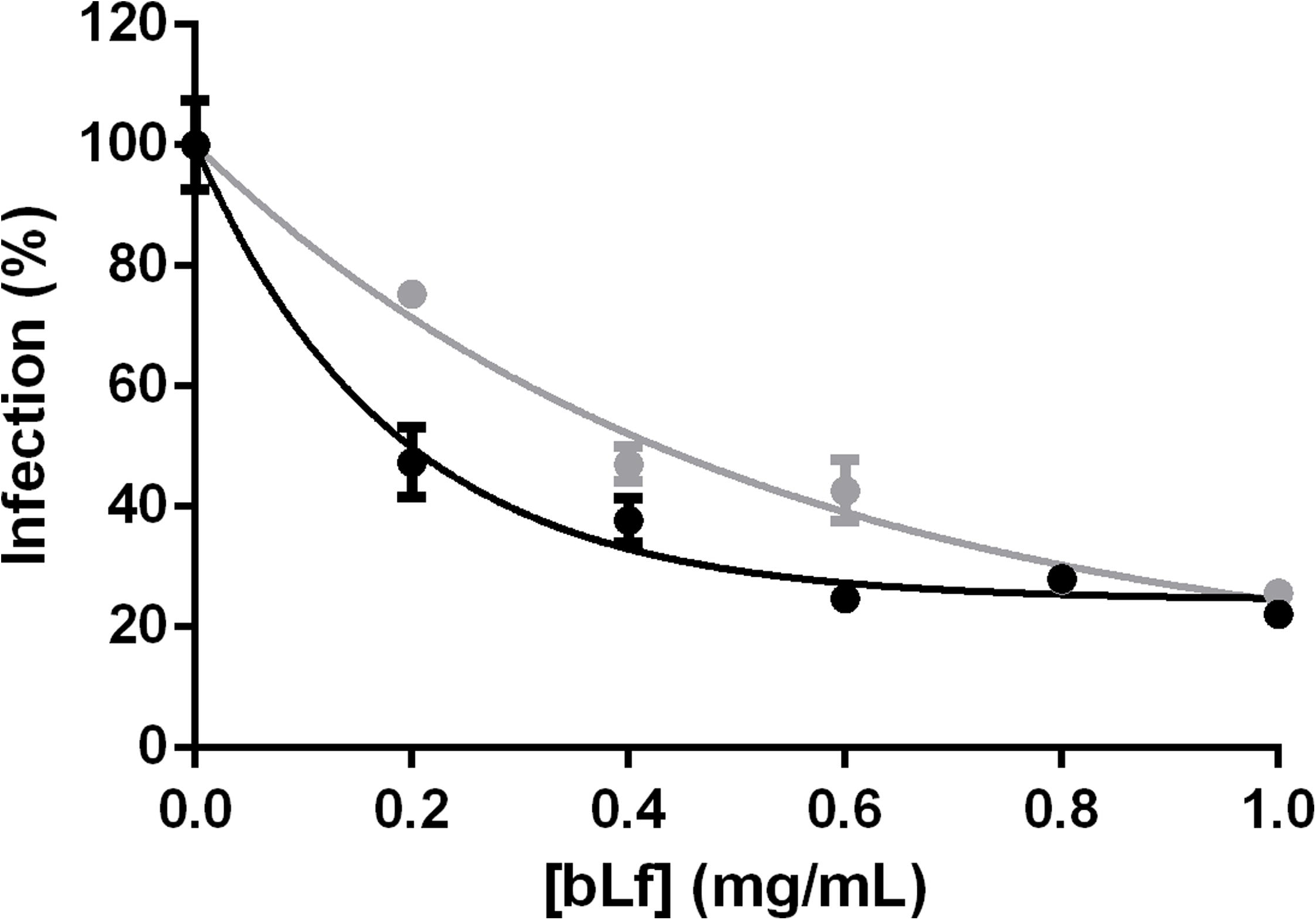
Dose-dependent inhibitory effect of bLf on CHIKV or ZIKV infection. Monolayers of Vero cells were incubated with the indicated concentrations of bLf at 37 °C throughout the course of infection by CHIKV (black) or ZIKV (gray), including immediately before (for 1 h), during (for 1 h), and immediately after (for 48 or 96 h, respectively) virus addition under the same MOI. Cells were stained and plaques were counted to determine the efficiency of infection. Data were obtained from 3 experiments and plotted as mean ± SD along with their exponential fittings, which revealed bLf IC_50_ values of 0.2 ± 0.005 mg/mL for CHIKV and 0.4 ± 0.006 mg/mL for ZIKV. All differences compared to the respective controls were extremely significant (P < 0.001).

### Inhibitory Effect of BLf on Different Stages of CHIKV or ZIKV Infection

A time-of-addition assay was next performed to determine the steps in CHIKV or ZIKV infection inhibited by bLf. In this approach, 1.0 mg/mL bLf was incubated with Vero cells before, during or after virus addition, and then tested as above for its effects on plaque formation. For both viruses, it was observed a significant antiviral activity of bLf at two of the three time points tested – before or during virus addition for CHIKV and during or after virus addition for ZIKV (Fig 3). Nevertheless, this inhibitory effect was clearly more pronounced when the protein was present together with the viruses, preventing CHIKV infection by approximately 70% and ZIKV infection by approximately 75%. When bLf was present before virus addition, it significantly inhibited CHIKV (approximate inhibition of 25%) but not ZIKV; inversely, when the protein was present after virus addition, it significantly inhibited ZIKV (approximate inhibition of 60%) but not CHIKV. Despite the large inhibitory effect promoted by bLf when it was present together with the viruses, CHIKV or ZIKV pretreatment with 1.0 mg/mL bLf for 1 h at 4 °C showed no significant deleterious effects on virus infectious titers (data not shown).

**Fig 3.**
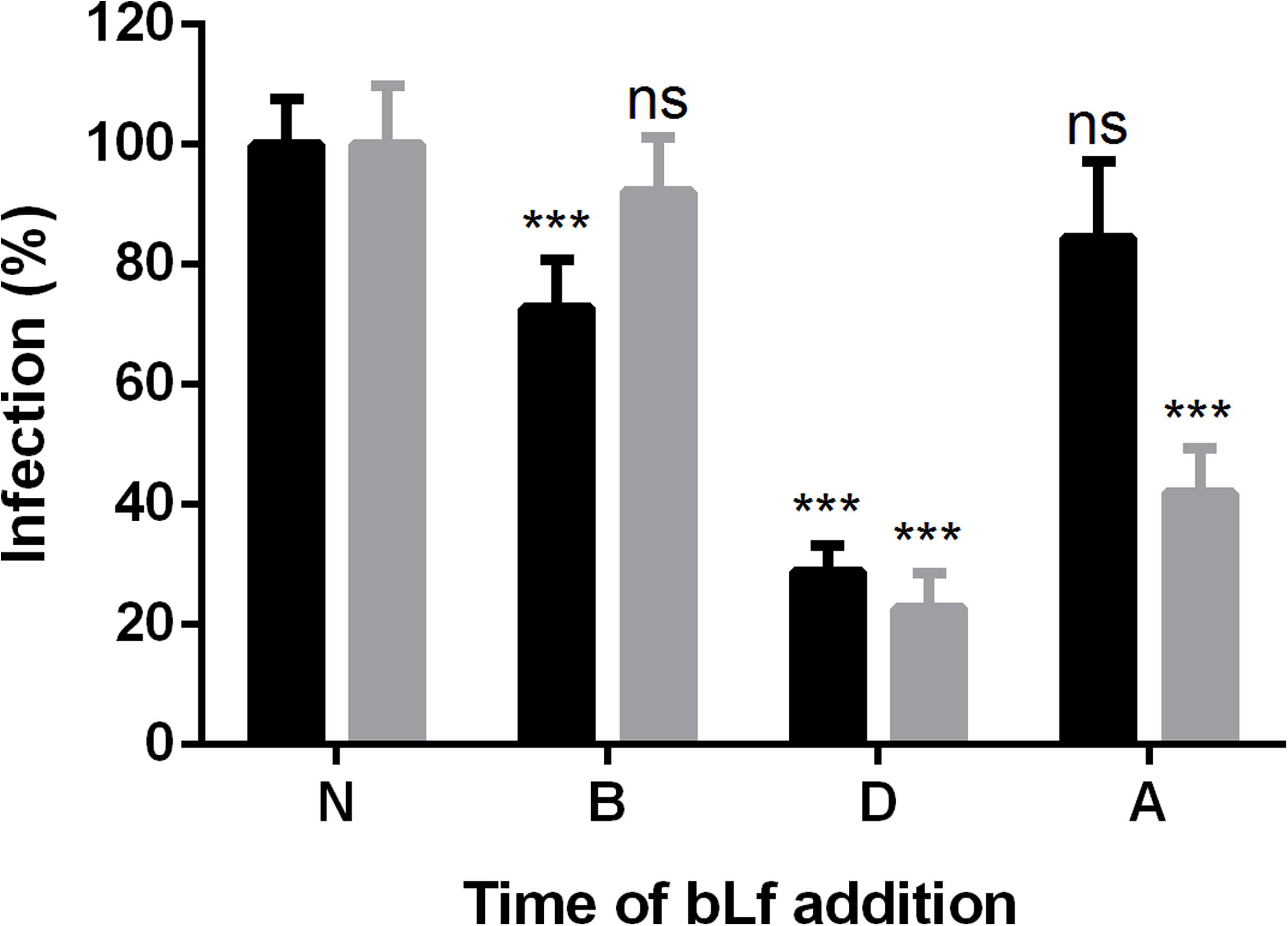
Inhibitory effect of bLf on different stages of CHIKV or ZIKV infection. Monolayers of Vero cells infected with CHIKV (black) or ZIKV (gray) under the same MOI were treated with 1.0 mg/mL bLf at different steps of the infection procedure: (N) never, (B) before virus addition, (D) during virus addition, or (A) after virus addition. At 48 h post-infection for CHIKV or 96 h postinfection for ZIKV, cells were stained and plaques were counted to determine the efficiency of infection. Data were obtained from 5 experiments and plotted as mean ± SD. Differences compared to the respective controls were either not significant (ns, P ≥ 0.05) or extremely significant (***, P<0.001).

### Anti-CHIKV/ZIKV Activity of BLf at the Level of Virus Binding

Since it seemed clear that bLf was mostly inhibiting an early event in the virus life cycle, the protein was tested for its ability to prevent virus infection by interfering with virus binding to the cell surface. In this experiment, Vero cells were first treated with 1.0 mg/mL bLf at 4 °C to retain protein molecules at the cell surface and then briefly incubated with CHIKV or ZIKV at the same temperature after washing away unbound protein molecules, being afterwards washed again to remove unbound virus particles and incubated at 37 °C to allow for infection progress. As assessed by iIFA, bLf-treated cells showed very low levels of infection for both viruses when compared to mock-treated cells (Fig 4).

**Fig 4.**
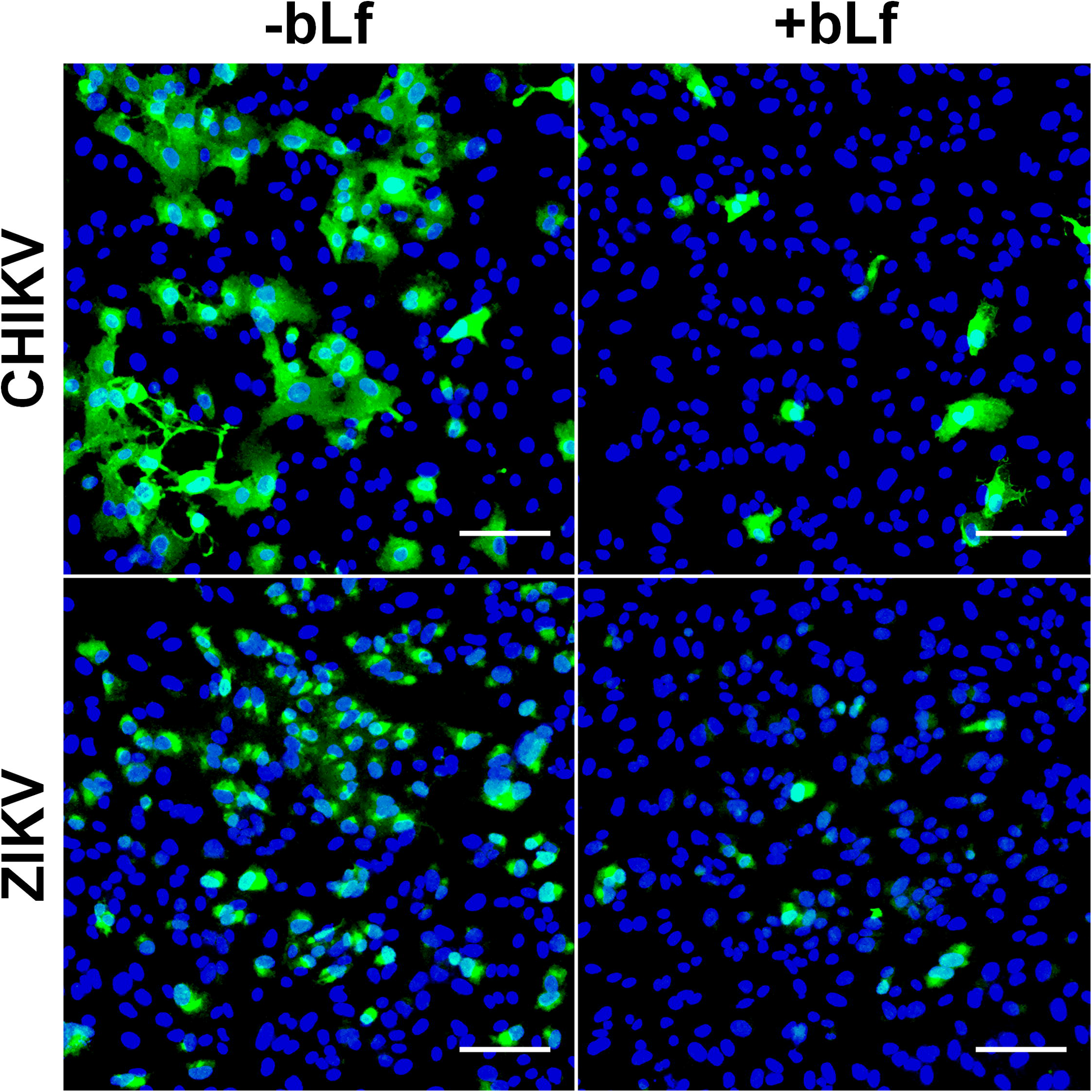
Anti-CHIKV/ZIKV activity of bLf at the level of virus binding. Monolayers of Vero cells were mock-treated (-bLf) or treated with 1.0 mg/mL bLf (+bLf) for 15 min at 4 °C, washed to remove unbound protein molecules, and then incubated with CHIKV (up) or ZIKV (down) under an MOI of 1 PFU/cell for another 15 min at 4 °C, being afterwards washed again to remove unbound virus particles and incubated at 37 °C to allow for infection progress. At 24 h post-infection for CHIKV or 48 h post-infection for ZIKV, cells were subjected to iIFA with anti-CHIKV or anti-ZIKV primary mouse polyclonal antibodies and FITC-conjugated anti-mouse IgG secondary goat polyclonal antibodies (green), in addition to nuclear staining with Hoechst 33342 (blue). Images were collected from 8 random visual fields and representative fluorescence channels of both probes were merged into a single channel for every experimental condition. Scale bars: 100 µm.

### Anti-CHIKV/ZIKV Activity of BLf at the Level of Virus Replication

To further investigate the relatively minor antiviral effects of bLf exerted at a post-entry step in virus infection, the protein was tested for its ability to reduce virus production by interfering with virus replication inside the cell. In this experiment, Vero cells were first incubated with CHIKV or ZIKV at 37 °C to allow for the entry of virus particles into the cell and then briefly treated with 1.0 mg/mL bLf at the same temperature after washing away unbound virus particles, being afterwards washed again to remove unbound protein molecules and incubated at 37 °C to allow for infection progress. As assessed by qRT-PCR, the supernatant of the bLf-treated cell culture showed approximately half of the virus load for both viruses when compared to the supernatant of the mock-treated cell culture (Fig 5).

**Fig 5.**
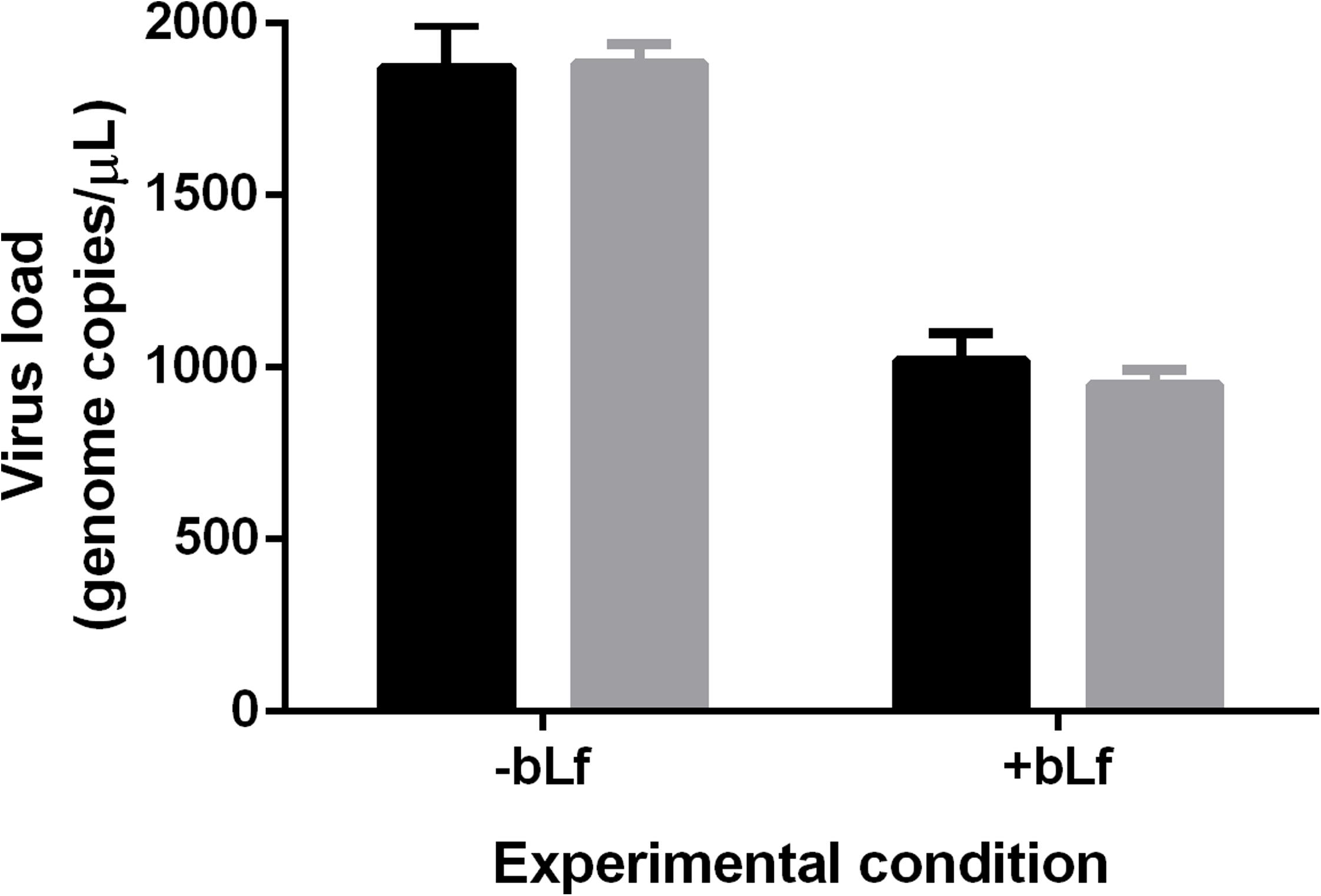
Anti-CHIKV/ZIKV activity of bLf at the level of virus replication. Monolayers of Vero cells were incubated with CHIKV (black) or ZIKV (gray) under an MOI of 0.1 PFU/cell for 1 h at 37 °C, washed to remove unbound virus particles, and then mock-treated (-bLf) or treated with 1.0 mg/mL bLf (+bLf) for another 1 h at 37 °C, being afterwards washed again to remove unbound protein molecules and incubated at 37 °C to allow for infection progress. At 24 h post-infection for CHIKV or 48 h post-infection for ZIKV, cell culture supernatants were subjected to RNA isolation followed by qRT-PCR with specific primers against defined sequences in CHIKV or ZIKV genomes. Data were obtained from 3 experiments and plotted as mean ± SD. Differences between respective +bLf and -bLf conditions were extremely significant (P<0.001).

## Discussion

Despite much in evidence, CHIKV and ZIKV are just the tip of the iceberg when it comes to the myriad of arboviruses that may emerge or reemerge in tropical and temperate regions of the world, specially in the Americas [20]. However, no selective inhibitors are available for a multitude of (re)emerging medically important viruses – in this scenario, broad-spectrum antiviral agents such as bLf, may offer important clues to cope with the challenge [21].

This study investigated whether the antiviral properties of bLf may be extended to CHIKV or ZIKV infection in Vero cells. Our data revealed a dose-dependent strong inhibitory effect by the protein in both cases, with no significant cytotoxicity, reaching a similar maximum inhibition of nearly 80% at 1.0 mg/mL via different IC_50_ values (∼0.2 mg/mL for CHIKV and ∼0.4 mg/mL for ZIKV). Previous studies using bLf against a different emerging alpha – Mayaro virus (MAYV) – or flavivirus – Japanese Encephalitis virus (JEV), demonstrated higher IC_50_ values (∼0.4 mg/mL and ∼0.5 mg/mL, respectively) in comparison to the respective virus counterparts addressed in this work [16, 22]. Such a difference indicates that CHIKV and ZIKV are even more sensitive than MAYV and JEV, respectively, to the effects of bLf.

The inhibitory activity of bLf over CHIKV or ZIKV infection was mostly exerted at a pre-entry step in virus infection (presumably binding), but the protein also affected a post-entry step in this process (presumably replication). It is worth noting that these observations, derived from the iIFA and qRT-PCR experiments, are not in contradiction with their counterparts derived from the timeof-addition experiment, as the analyses were performed under slightly different conditions by approaches that assess virus infection efficiency from different standpoints.

While in the iIFA experiment bLf pretreatment was carried out at 4 °C, in the time-of-addition experiment this procedure was carried out at 37 °C. Since both endocytosis and vesicle trafficking are active at 37 °C but not at 4 °C, the occurrence of only a slight antiviral effect in the time-of-addition experiment is probably associated with partial bLf internalization and fast glycosaminoglycan turnover, to which the protein is known to bind [23]. Regarding the comparison between the observations derived from the qRT-PCR and the time-of-addition experiments, it is important to bear in mind that post-entry events which only partially impair the virus infection process not necessarily lead to plaque number reduction, since even a minimal amount of virus progeny is able to promote the radial death zone that characterizes the plaque. Thus, as virus replication was not fully inhibited when bLf was added after virus entry, some virus progeny was still able to be produced and account for plaque formation in the time-of-addition experiment.

Previous studies explain the common effects of Lf on virus binding by the blockage of cell-surface glycosaminoglycans such as heparan-sulfate, exploited by many virus species as an inespecific adhesion molecule, while the rare effects of Lf on virus replication are explained by the induction of interferon (IFN)-α/β antiviral cytokine expression [24]. Although bLf has nearly 70% amino acid sequence identity with hLf [25], the bovine version of the protein is often reported to exhibit higher antiviral activity than its human version [26]. Moreover, iron-unsaturated Lf (apoLf) is more potent than its iron-saturated isoform (holoLf) against some virus species [27]. Interestingly, Lf also contains various conserved peptides which are released upon its hydrolysis by proteases and still retain the antimicrobial activity [28].

The risk of CHIKV and ZIKV adaptation to urban mosquito vectors other than *Aedes aegypti* and *Aedes albopictus* – such as *Culex quinquefasciatus* – due to the current rampant spreading of these viruses, specially in Latin America, may predict an even greater geographical dispersion of their respective diseases [29]. Added to this, the risk of CHIKV/ZIKV introduction in a new sylvatic environment – such as the Amazon rainforest – may establish permanent virus reservoirs for constant outbreaks in the newly-affected areas, similar to the sylvatic cycle of yellow fever in Brazil [30]. Given the current scenario and these potential risks, there is an urgency for efficient prophylactic and therapeutic approaches against Chikungunya and Zika fevers. Our work shows that the antiviral properties of bLf are extendable to CHIKV and ZIKV and may be explored to design a two-in-one strategy against their infections.

## Acknowledgments

We thank the technical staff of Seção de Arbovirologia e Febres Hemorrágicas, Instituto Evandro Chagas, Ministério da Saúde, for competent assistance.

## References

1. Robinson MC. An epidemic of virus disease in Southern Province, Tanganyika Territory, in 1952-53. I. Clinical features. Trans R Soc Trop Med Hyg 1955; 49:28–32.

2. Lumsden WHR. An epidemic of virus disease in Southern Province, Tanganyika Territory, in 1952-53. II. General description and epidemiology. Trans R Soc Trop Med Hyg 1955; 49:33–57.

3. Dick GWA, Kitchen SF, Haddow AJ. Zika virus. I. Isolations and serological specificity. Trans R Soc Trop Med Hyg 1952; 46:509–20.

4. Dick GWA. Zika virus. II. Pathogenicity and physical properties. Trans R Soc Trop Med Hyg 1952; 46:521–34.

5. Cardoso CW, Paploski IAD, Kikuti M, et al. Outbreak of exanthematous illness associated with Zika, Chikungunya, and Dengue viruses, Salvador, Brazil. Emerg Infect Dis 2015; 21:2274–6.

6. Fauci AS, Morens DM. Zika virus in the Americas – Yet another arbovirus threat. N Engl J Med 2016; 374:601–4.

7. Shan C, Xie X, Barrett ADT, et al. Zika virus: diagnosis, therapeutics, and vaccine. ACS Infect Dis 2016; 2:170–2.

8. Martinez JP, Sasse F, Brönstrup M, Diez J, Meyerhans A. Antiviral drug discovery: broadspectrum drugs from nature. Nat Prod Rep 2015; 32:29–48.

9. Baker EN, Baker HM. Molecular structure, binding properties and dynamics of lactoferrin. Cell Mol Life Sci 2005; 62:2531–9.

10. Groves ML. The isolation of a red protein from milk. J Am Chem Soc 1960; 82:3345–50.

11. Johanson B. Isolation of an iron-containing red protein from human milk. Acta Chem Scand 1960; 14:510–2.

12. Masson PL, Heremans JF, Dive CH. An iron-binding protein common to many external secretions. Clin Chim Acta 1966; 14:735–9.

13. Masson PL, Heremans JF, Schonne E. Lactoferrin, an iron-binding protein in neutrophilic leukocytes. J Exp Med 1969; 130:643–58.

14. Jenssen H, Hancock REW. Antimicrobial properties of lactoferrin. Biochimie 2009; 91:19–29.

15. Dulbecco R, Vogt M. Some problems of animal virology as studied by the plaque technique. Cold Spring Harb Symp Quant Biol 1953; 18:273–9.

16. Carvalho CAM, Sousa Jr IP, Silva JL, Oliveira AC, Gonçalves RB, Gomes AMO. Inhibition of Mayaro virus infection by bovine lactoferrin. Virology 2014; 452-453:297–302.

17. Tikasingh ES, Spence L, Downs WG. The use of adjuvant and sarcoma 180 cells in the production of mouse hyperimmune ascitic fluids to arboviruses. Am J Trop Med Hyg 1966; 15:219–26.

18. Lanciotti RS, Kosoy OL, Laven JJ, et al. Chikungunya virus in US travelers returning from India, 2006. Emerg Infect Dis 2007; 13:764–7.

19. Lanciotti RS, Kosoy OL, Laven JJ, et al. Genetic and serologic properties of Zika virus associated with an epidemic, Yap State, Micronesia, 2007. Emerg Infect Dis 2008; 14:1232–9.

20. Vasconcelos PFC, Calisher CH. Emergence of human arboviral diseases in the Americas, 2000-2016. Vector Borne Zoonotic Dis 2016; 16:295–301.

21. Debing Y, Neyts J, Delang L. The future of antivirals: broad-spectrum inhibitors. Curr Opin Infect Dis 2015; 28:596–602.

22. Chien YJ, Chen WJ, Hsu WL, Chiou SS. Bovine lactoferrin inhibits Japanese encephalitis virus by binding to heparan sulfate and receptor for low density lipoprotein. Virology 2008; 379:143–51.

23. Mann DM, Romm E, Migliorini M. Delineation of the glycosaminoglycan-binding site in the human inflammatory response protein lactoferrin. J Biol Chem 1994; 269:23661–7.

24. Wakabayashi H, Oda H, Yamauchi K, Abe F. Lactoferrin for prevention of common viral infections. J Infect Chemother 2014; 20:666–71.

25. Pierce A, Colavizza D, Benaissa M, et al. Molecular cloning and sequence analysis of bovine lactotransferrin. Eur J Biochem 1991; 196:177–84.

26. Berlutti F, Pantanella F, Natalizi T, et al. Antiviral properties of lactoferrin – a natural immunity molecule. Molecules 2011; 16:6992–7018.

27. Van der Strate BWA, Beljaars L, Molema G, Harmsen MC, Meijer DKF. Antiviral activities of lactoferrin. Antiviral Res 2001; 52:225–39.

28. Sinha M, Kaushik S, Kaur P, Sharma S, Singh TP. Antimicrobial lactoferrin peptides: the hidden players in the protective function of a multifunctional protein. Int J Pept 2013; doi: 10.1155/2013/390230.

29. Ayres CF. Identification of Zika virus vectors and implications for control. Lancet Infect Dis 2016; 16:278–9.

30. Favoretto S, Araújo D, Oliveira D, et al. First detection of Zika virus in neotropical primates in Brazil: a possible new reservoir. BioRxiv 2016; doi: 10.1101/049395.

